# A Stochastic Programming Approach to the Antibiotics Time Machine Problem

**DOI:** 10.1101/2023.08.31.555704

**Authors:** Oğuz Mesüm, Ali Rana Atilgan, Burak Kocuk

## Abstract

Antibiotics Time Machine is an important problem to understand antibiotic resistance and how it can be reversed. Mathematically, it can be modelled as follows: Consider a set of genotypes, each of which contain a set of mutated and unmutated genes. Suppose that a set of growth rate measurements of each genotype under a set of antibiotics are given. The transition probabilities of a ‘realization’ of a Markov chain associated with each arc under each antibiotic are computable via a predefined function given the growth rate realizations. The aim is to maximize the expected probability of reaching to the genotype with all unmutated genes given the initial genotype in a predetermined number of transitions, considering the following two sources of uncertainties: i) the randomness in growth rates, ii) the randomness in transition probabilities, which are functions of growth rates. We develop stochastic mixed-integer linear programming and dynamic programming approaches to solve static and dynamic versions of the Antibiotics Time Machine Problem under the aforementioned uncertainties. We adapt a Sample Average Approximation approach that exploits the special structure of the problem and provide accurate solutions that perform very well in an out-of-sample analysis.

## 1 Introduction

Antimicrobial resistance is estimated to have played a role in 4.95 million deaths in 2019 [1]. To combat antibiotic resistance (ABR), we traditionally have confined our attention to discover and design new antibiotics [2]. In addition to the utilization of artificial intelligence in these efforts [3, 4], two new lines of research have recently emerged: (i) Developers have repurposed already available drugs against ABR [5, 6], or (ii) researchers have prescribed a sequence of antibiotics that exploits the collateral sensitivity of bacterial mutations to slow down or reverse the resistance [7, 8]. Even though there were a limited number of clinical observations that raised opposing views [9, 10], later experimental and computational studies suggested that there might still exist optimal drug ordering procedures to delimit ABR [11, 12, 13, 14, 15, 16, 17], including the recently developed reinforcement learning approach [18]. An adaptive methodology substituting a pre-determined schedule, with one based on a “dynamic” recipe in which the drugs are selected while the treatment is in progress, continues to be a pressing need.

Here is a recipe that takes advantage of the myopic drug response of bacteria: Suppose we have a population of bacteria and for each bacterium of a given genotype, an antibiotic alters it to another type with some probability. For a viable drug policy, we apply a number of antibiotics in a specified sequence so as to maximize the fraction of bacteria reverting to the wild type. A straightforward approach would be to explicitly enumerate all the permutations with repetitions, and then compute the probability of returning to the wild type [7]. Such an explicit enumeration algorithm is prohibitively time-consuming, even for moderate sized problems. Nichol et al. [8] developed a weighted random walk through the fitness landscape defined by a drug; accordingly, they can navigate the genotype space to avoid ABR. This simulation methodology was later repeated and supported by extensive number of experimental constructs [12, 13, 14]. Tran and Yang [19] proved that an optimized walk that is initiated at any point of the rugged landscape and terminated at the station of the wild genotype, referred to as an “antibiotic time machine”, is NP-hard to compute. A mixed-integer linear programming (MILP) reformulation, which is a widely used approach in the optimization community with several efficient computational tools, was proposed in [17] to solve this problem. In fact, this latter approach turned out to be orders-of-magnitude faster than complete enumeration.

In our paper, we will refer to the models in the line of research described above as “static” and “deterministic” (we will make these notions more precise below), and extend it from two aspects: Firstly, these recent efforts concerning the Antibiotics Time Machine Problem has only considered a “static” version of the problem in which all the antibiotics administration decisions are made in the beginning and executed without observing the intermediate random outcomes. In our paper, we develop a “dynamic” version of the problem in the sense that the next antibiotic administration decision is made by observing the realizations of the previous such actions. We solve the resulting problem using another well-developed technique from the optimization literature called Dynamic Programming (DP), which has recently been applied to cancer therapy [20]. Clearly, this has the potential to increase the probability of reversing the resistance. Secondly, the literature on the Antibiotics Time Machine Problem have only considered the randomness associated with administering a certain antibiotic. Typically, this randomness is modelled as a Markov chain where the transition probabilities are a function of the experimentally obtained “growth rates” and the corresponding optimization problem can be modelled as deterministic. However, if these growth rates are measured multiple times, which is typically the case (see, e.g. [21]), an additional randomness due to these growth rate measurements are observed. To the best of our knowledge, our paper is the first study that considers both the randomness in growth rates and the transition probabilities. We formulate a “stochastic” version of the Antibiotics Time Machine Problem as a stochastic optimization problem. A significant challenge in this optimization problem is the computational effort of computing the transition probability matrices, for which we use the well-known optimization technique called the Sample Average Approximation (SAA).

We conduct extensive computational experiments using the four-allele real dataset from [21] with 23 antibiotics under two probability models from [7]. The results reveal that our SAA-based stochastic optimization models are extraordinarily accurate as the in-sample and out-of-sample values for the probability of reaching the wild type are at most 0.01 away from each other. Under the static stochastic model, we report that the probability of reaching the wild type is at least 0.495 and 0.695 for two probability models under the static model with a treatment length of 16 for all different initial genotypes. On the other hand, under the dynamic stochastic model, this probability already reaches 0.995 for treatments plans of length 11 and 14. We believe that these remarkable results suggest that optimization techniques can be instrumental in solving various versions of the Antibiotic Time Machine Problem.

## 2 Methods

**Notation 1**. We use bold letters to denote vectors and matrices. We assume that all vectors are column vectors. We use the notation ***e***_***i***_ for the *i*-th unit vector, ***e*** for the vector of ones, and 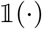 for the indicator function. We also use the convention 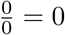.

### 2.1 Deterministic Model

In this section, we consider a deterministic model for the Antibiotics Time Machine Problem in the sense that the transition probability matrices governing the effect of antibiotics are precisely known and given.

#### 2.1.1 Problem Definition

Suppose that we are interested in a fitness landscape with *a* many alleles. We will denote a genotype by a vector ***g***. Each allele *h ∈* {1, …, *a*} in ***g*** can either be mutated (*gh* = 1) or unmutated (*g*_*h*_ = 0). Consequently, the total number of genotypes in a fitness landscape, denoted by *d*, with *a* many alleles will be 2^*a*^. We will call the special genotype **0** as the *wild type*. Suppose that each genotype ***g*** ∈ {0, 1}^*a*^ has a unique ID^1^ from the set {1, …, *d*}.

Suppose that we have *K* antibiotics. The effect of an antibiotic is modelled as a Markov chain with a transition probability matrix ***M***^***k***^ **∈** ℝ^*d×d*^, *k* = 1, …, *K*. In other words, when antibiotic *k* is applied to a genotype with ID *i*, then the next genotype ID becomes *j* with probability 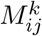. Due to the widely accepted assumption of Gillespie [22, 23] called *Strong Selection Weak Mutation*, the transitions between two genotypes ***g*** and ***g***^***′***^ can occur only if a single allele is different, that is, ‖***g*** *−* ***g***^***′***^‖_1_ = 1. Let us denote the set of such pairs as *J* .

The aim in the Antibiotics Time Machine Problem is to come up with a plan of predetermined number of antibiotic intakes that maximizes the probability of reaching the wild type starting from a given initial genotype. Depending on the type of the plans allowed, we will consider two versions of the problem: Static and Dynamic. In the Static Version, we decide the sequence of antibiotics to be applied in the beginning of the planning horizon whereas in the Dynamic Version, we come up with a policy telling us which antibiotic to take at each decision epoch based on the current genotype. Observe that the Dynamic Version requires intermediate observations regarding the state transitions to be observable. We formally state these two versions as below:

**Problem 1** (Static Deterministic Version). Given *K* antibiotics with their transition probability matrices ***M***^***k***^, *k* = 1, …, *K*, the length of the treatment plan *N* and the initial distribution ***p***, determine a *sequence* of antibiotics to be applied so that the probability of reaching the desired final distribution ***q*** is maximized.

**Problem 2** (Dynamic Deterministic Version). Given *K* antibiotics with their transition probability matrices ***M***^***k***^, *k* = 1, …, *K*, the length of the treatment plan *N* and the initial distribution ***p***, determine a *policy* of antibiotics to be applied so that the probability of reaching the desired final distribution ***q*** is maximized.

In general, the initial distribution vector ***p*** is the unit vector 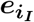 corresponding to the initial genotype *iI*, and the desired final distribution vector ***q*** is the unit vector corresponding to the wild type although most of our solution approaches allow for a general initial and final distributions.

**Example 1**. Let us now clarify the problem setting with a toy example using Figure 1. In this example, we have *a* = 2 alleles and *d* = 2^*a*^ = 4 genotypes. The nodes correspond to the genotypes and the numbers near each arc are the transition probabilities. For now, let us assume that these numbers are given. We will explain how to obtain these probabilities using the growth rates, which are given with numbers in parenthesis above each node, in Section 2.1.2.

**Figure 1:**
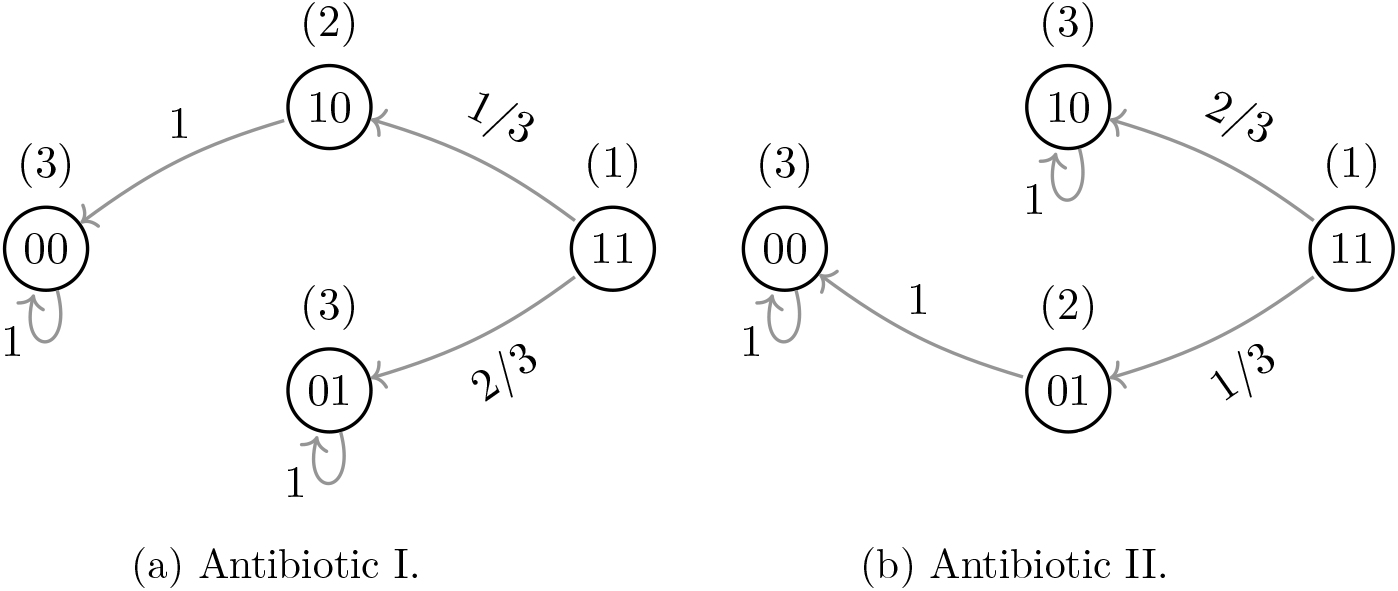
Markov chains corresponding to two antibiotics.

Suppose that the initial genotype is 11 and our aim is to maximize the probability of reaching the wild type in *N* = 2 steps. In this case, there are four possible sequences for Problem 1: I-I, I-II, II-I, II-II with respective probabilities of 1*/*3, 2*/*3, 2*/*3, 1*/*3. Therefore, we deduce that the optimal sequences are I-II and II-I. On the other hand, there are eight possible policies for Problem 2: I-(I,I), I-(I,II), I-(II,I), I-(II,II), II-(I,I), II-(I,II), II-(II,I), II-(II,II). Here, in a policy given by the notation *y*_1,11_ *−* (*y*_2,10_, *y*_2,01_), *y*_1,11_ is the antibiotic applied in step 1, and antibiotics *y*_2,10_ and *y*_2,01_ are the ones applied in step 2 if the observed state is 10 and 01, respectively. It is easy to see that the optimal policies are I-(I,II) and II-(I,II) with the optimal value of 1. As expected, the added flexibility of observing the state transition after step 1 and making a decision accordingly increases the probability of reaching the wild type.

#### 2.1.2 Transition Probability Matrix Computation

An important aspect of the Antibiotics Time Machine Problem is the computation of the transition probability matrices, for which we use the framework from [7]. The authors measure the growth rate of each genotype under the administration of an antibiotic and store it in a vector 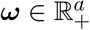. The rationale behind the computation of the matrices is the following: For a pair of neighbors (*j, j*^*′*^) ∈ *J*, the transition probability from genotype *j* to genotype *j*^*′*^ is positive if the growth rate of genotype *j*^*′*^ is larger than the growth rate of genotype *j*, that is, *ω*_*j*_ *> ω*_*j*_; and zero otherwise. Following [7], we consider two models regarding how the precise transition probabilities are computed:

- Correlated Probability Model (CPM):

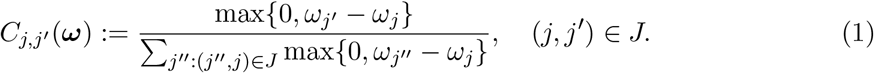

- Equal Probability Model (EPM):

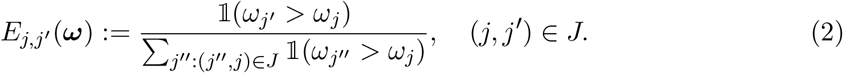

For each antibiotic *k*, we will denote its growth rate vector as ***ω***^***k***^, and set the transition probability matrix ***M***^***k***^ = ***C*(*ω***^***k***^) if CPM is used and ***M***^***k***^ = ***E*(*ω***^***k***^) if EPM is used.

In Example 1 illustrated with Figure 1, the numbers in paranthesis above each node give the growth rate of the corresponding genotype under each antibiotic. The transition probabilities are computed using CPM in this example. If EPM was utilized, the transitions would occur with equal probability at the genotype 11 for each antibiotic.

#### 2.1.3 Static Optimization

We now review Problem 1, that is, the Static Deterministic Version of the Antibiotics Time Machine Problem. Suppose that our aim to maximize the probability of reaching to the final distribution ***q*** starting from the initial distribution ***p*** in *N* steps using transition probability matrices from a finite set *ℳ* := *{****M***^***k***^ : *k* = 1, …, *K}*. Then, we consider

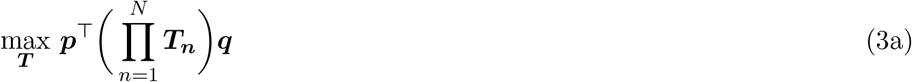

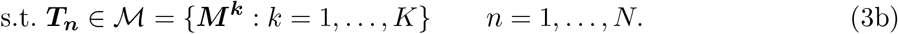

This formulation first appeared in [19] in which the authors show that Problem 1 is NP-Hard.

A straightforward approach to attack formulation (3) is to use complete enumeration, which requires computing the objective function for *K*^*N*^ possible orderings as proposed in [7]. However, this approach is computationally expensive as it only allows limited number of steps to be considered in a practical setting.

Recently, an MILP reformulation is proposed in [17] to solve Problem 1. The key idea of this approach is to define two sets of decisions variables: We will denote the first set of decision variables as *x*_*n,k*_ that takes value one if antibiotic *k* is applied in step *n* and zero otherwise. The second set of decision variables, denoted as 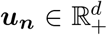, represent the probability distribution after *n* transitions.

Notice that these variables satisfy the recursion

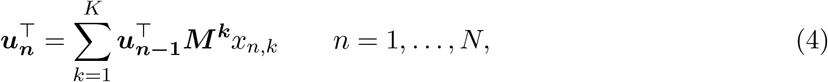

where ***u***_**0**_ = ***p***. Finally, relation (4) is linearized using a disjunctive formulation by defining the copy variables 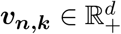. The precise MILP model is given as below:

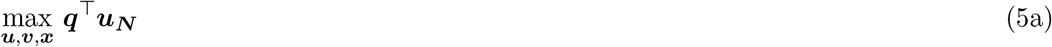

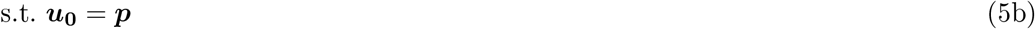

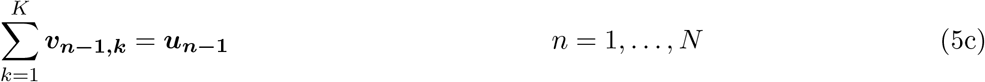

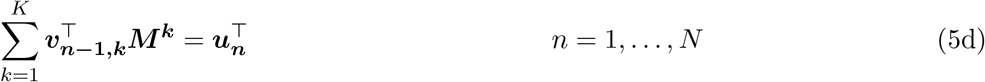

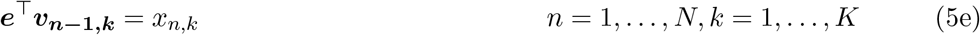

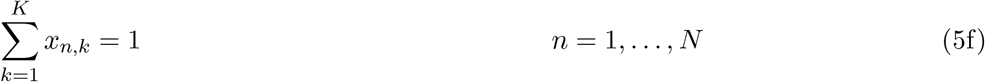

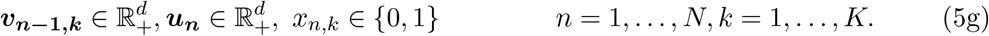

Here, the objective function (5a) maximizes the probability of reaching the desired distribution after *N* transitions. Constraints (5b)–(5d) linearize the relation (4) using a disjunctive argument. whereas constraint (5e) guarantees that one antibiotic is applied in each step *n*. We refer the reader to [17] for the details of this model and the computational evidence, which, expectedly, shows the superior performance of the MILP model compared to the complete enumeration.

#### 2.1.4 Dynamic Optimization

We now formally analyze Problem 2, that is, the Dynamic Deterministic Version of the Antibiotics Time Machine Problem. In particular, we propose a DP-based approach to solve this version. In this approach, we will assume that the initial and final distribution vectors ***p*** and ***q*** are unit vectors corresponding to states *i*_*I*_ and *i*_*F*_, respectively. Let us define *ν*_*n*_(*i*) as the maximum probability of reaching the final state *i*_*F*_ at the end of *N* -th transition given that we are at state *i* after *n* transitions. With this definition, we have the following boundary value:

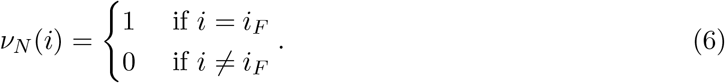

Solving the Dynamic Version of the Antibiotics Time Machine Problem amounts to computing *ν*_0_(*i*_*I*_).

Let *y*_*n,i*_ *∈ {*1, …, *K}* be the antibiotic applied in step *n* when the state is *i*. We can state the optimality conditions as follows:

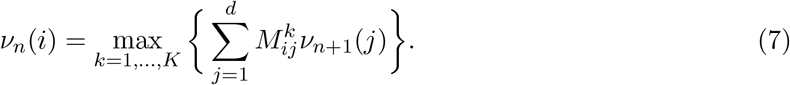

We then use a backward DP approach given in Algorithm 1 to solve Problem 2.

##### Algorithm 1 Backward DP.

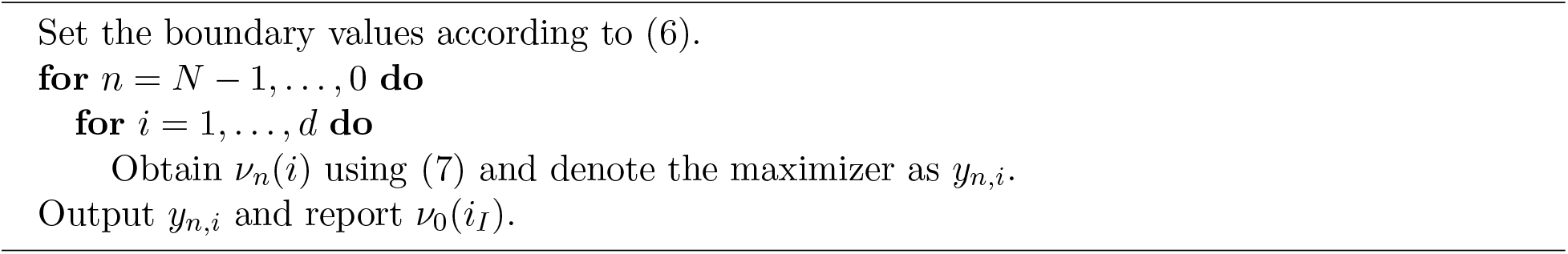

##### Proposition 1.

*Problem 2 is polynomially solvable in problem parameters d, K and N. Proof*. The statement holds true since the running time of Algorithm 1 is *O*(*d*^2^*KN*).

**Remark 1**. *It is possible to solve Problem 2 via an MILP as well. However, our preliminary experiments have shown that such as approach is extremely time consuming compared to the DP approach described in this section. Therefore, we have chosen not to use MILP for Problem 2*.

### 2.2 Stochastic Model

The analysis of the Antibiotics Time Machine Problem in Section 2.1 assumes that the transition probability matrices governing the Markov chains associated with each antibiotic are deterministic.

The underlying reason for this assumption is that the growth rates for each antibiotic are coming from a single measurement (see, e.g., [7]). However, in practice, the biologists do not rely on a single experiment to estimate important quantities such as growth rates, rather, they replicate their experiments under same ambient conditions (see, e.g., [21]). Even under same conditions, it is expected that the growth rates to change from one replication to another. Hence, it makes sense to model the growth rate of each genotype under the administration of an antibiotic as a random variable, which, in turn, causes randomness in the transition probability matrices. In this section, we consider a stochastic model for the Antibiotics Time Machine Problem in order to account for the stochasticity of the transition probability matrices governing the effect of antibiotics.

#### 2.2.1 Uncertainty Modeling and Problem Definition

In this subsection, we first explain the cause of the uncertainty, and then formally define the stochastic versions of the Antibiotics Time Machine Problem.

Suppose that growth rate measurements under each antibiotic are repeated *R* times. In particular, let the growth rate of genotype *j* under the administration of antibiotic *k* in replication *r* be given as 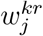. To give a perspective, the data acquired from [21] consists of the growth rates of *d* = 16 genotypes under *K* = 23 different antibiotics (or doses) measured for *R* = 12 times. We will assume that the *j*-th element of the growth rate vector 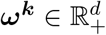 for antibiotic *k* is a random variable shown by 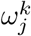, and it takes values from the set 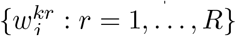 with equal probability.

As an example, consider the growth rate observations of antibiotic CEC retrieved from [21] as shown in Figure 2. Suppose that we apply this antibiotic to the genotype 1000. Clearly, even the smallest growth rate observation of this genotype is larger the largest growth rate observations of its neighbors 0000, 1100 and 1010, hence, no transitions will occur to these genotypes. However, a transition to genotype 1001 is possible but not certain.

**Figure 2:**
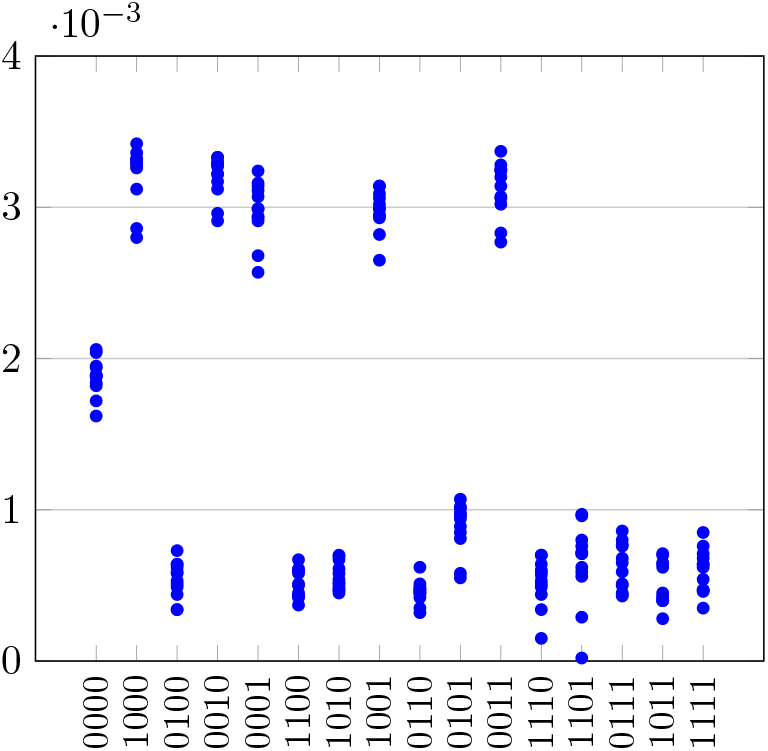
Growth rate observations of Cefaclor (CEC) with 4 μg*/*mL concentration [21].

Notice that if we decide to apply a certain antibiotic, any of *R*^*d*^ many transition probability matrices have an equal probability to be observed. Therefore, we need to come up with an approach that considers this aspect of the problem. To formalize the problem setting, we first define an index set *S* := *{*1, …, *R}*^*d*^. For any antibiotic *k* = 1, …, *K* and ***s*** *∈ S*, we define a vector ***ω***^***k***,***s***^ ∈ ℝ^*d*^ such that 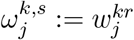 if *s*_*j*_ = *r* for *j* = 1, …, *d*. We will denote the matrix for an antibiotic *k* under the sample ***s*** as ***M***^***k***,***s***^, which is computed via *C*(***ω***^***k***,***s***^) or *E*(***ω***^***k***,***s***^) using formulas (1) or (2) depending on whether CPM or EPM is used.

**Example 1** (continuing from p. 4). Now, suppose that the growth rate measurements are replicated for both antibiotics and there are only two changes reported: The growth rate of genotype 00 under antibiotic I increases to 3 + *E* and the growth rate of genotype 00 under antibiotic II decreases to 3*−E*, for some small *E >* 0. The Markov chains corresponding to this replication is given in Figure 3.

**Figure 3:**
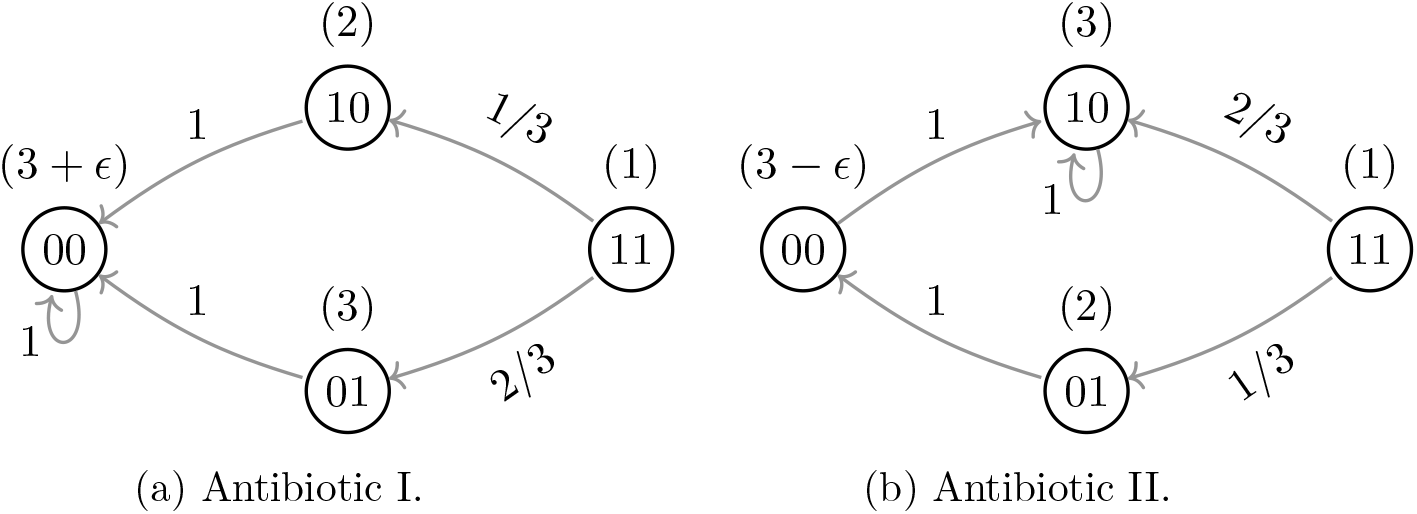
Markov chains corresponding to two antibiotics in Replication 2.

Solely based on the growth rate measurements from Replication 2, the optimal sequences for Problem 1 are I-I and II-I, and the optimal policies for Problem 2 are I-(I,I), I-(I,II), II-(I,I), II-(I,II) with optimal value of 1.

We are now ready to formally define two stochastic versions of the Antibiotics Time Machine Problem.

**Problem 3** (Static Stochastic Version). Given *K* antibiotics with their uncertain but equally likely transition probability matrices *{****M***^***k***,***s***^ : *s ∈ S}, k* = 1, …, *K*, the length of the treatment plan *N* and the initial distribution ***p***, determine a *sequence* of antibiotics to be applied so that the *expected* probability of reaching the desired final distribution ***q*** is maximized.

**Problem 4** (Dynamic Stochastic Version). Given *K* antibiotics with their uncertain but equally likely transition probability matrices *{****M***^***k***,***s***^ : *s ∈ S}, k* = 1, …, *K*, the length of the treatment plan *N* and the initial distribution ***p***, determine a *policy* of antibiotics so that the *expected* probability of reaching the desired final distribution ***q*** is maximized.

**Example 1** (continuing from p. 8). To solve Problem 3, we can compute the expected probability of reaching the wild type under each of the four sequences where the expectation is taken over the replications. In this case, the optimal sequence turns out to be II-I with the optimal value of 5/6. Similarly, to solve Problem 4, we can compute the expected probability of reaching the wild type under each of the eight policies. In this case, the optimal policies are I-(I,II) and II-(I,II) with the optimal value of 1.

**Remark 2**. For Example 1, it is interesting to note that the optimal solutions of Problems 3 and 4 can be obtained by solving them as particular instances of Problems 1 and 2 in which the *average transition probabilities* are used. In particular, consider Figure 4, which is obtained by *averaging out* Figures 1 and 3. The optimal solutions of Problems 3 and 4 for Example 1 considering two replications coincide with the optimal solutions of Problems 1 and 2 considering the average Markov chains from Figure 4, respectively. In fact, this is not a coincidence as we later show in Propositions 2 and 3.

**Figure 4:**
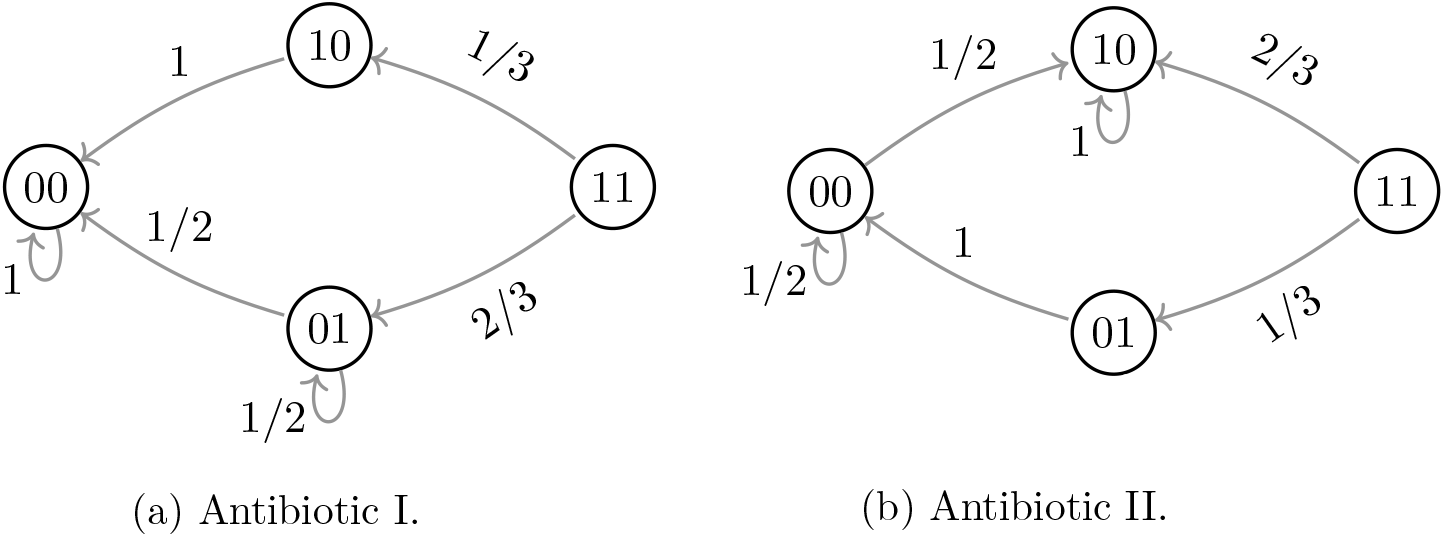
Average Markov chains corresponding to two antibiotics from two replications.

#### 2.2.2 Static Optimization

**Formulation** In this section, we will consider uncertainty in the transition probability matrices ***M***^***k***^. Recall that the randomness in growth rate observations results in randomness in the elements of the transition probability matrices. We assume that the growth rate observations from the replications are independent and identically distributed.

To capture the randomness in transition probability matrices, we change the optimization formulation (3) by the following model:

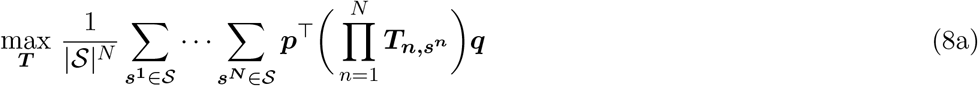

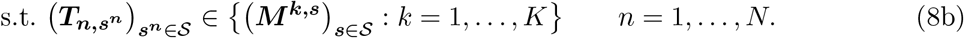

The main idea behind this optimization model is as follows: If we decide to apply antibiotic *k* in step *n*, then the transition probability matrices ***T***_***n***,***s***_***n*** take the value ***M***^***k***,***s***^ for each ***s*** *∈ S* (this is guaranteed through constraint (8b)). Then, the objective function (8a) computes the *expected* maximum probability of reaching to the final distribution ***q*** starting from the initial distribution ***p*** in *N* steps by considering all possible combinations of random transition probability matrices across each step.

Upon first glance, the optimization problem (8) looks considerably more difficult to solve than the optimization problem (3) due to the increased number of variables and constraints. However, we now prove that problem (8) can be formulated as an instance of problem (3).

**Notation 2**. We will denote the *average transition probability matrix for antibiotic k* by 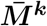, and compute it as

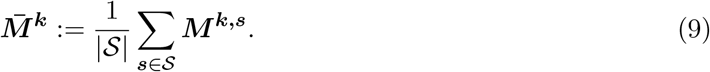

##### Proposition 2.

*Problem 3 is an instance of Problem 1*.

*Proof*. To prove this statement, we first note the following relation:

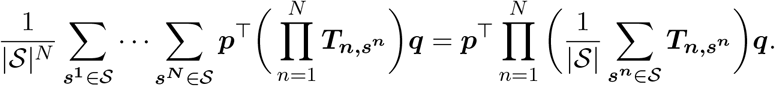

In addition, we observe that

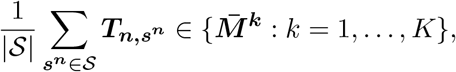

where 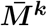 matrices are defined in (9). Therefore, Problem 3 reduces to an instance of Problem 1 with 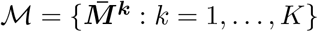, and can be solved by the MILP model (5).

Although Proposition 2 suggests a practical way to solve Problem 3, this is only possible if the average transition probability matrices 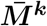 are available. Unfortunately, the complexity of computing each of these matrices is *O*(*R*^*d*^). To circumvent this issue, we propose a sampling approach to estimate the average transition probability matrices below.

##### Sample Average Approximation

Since the enumeration of all possible transition probability matrices is not very practical, we now propose a way to estimate the average transition probability matrix as follows. Let *S ⊆ S* be a randomly selected sample index set. Then, we compute the following estimator:

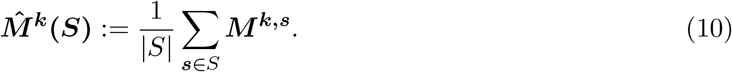

Since obtaining the set of matrices 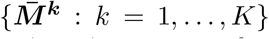 is computationally expensive, we will use Sample Average Approximation (SAA), see e.g. [24, 25, 26] to solve model (5) with the estimated matrices. In particular, given samples 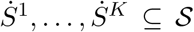, we solve model (5) with 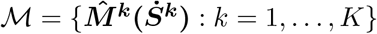. Suppose that the objective function is computed as

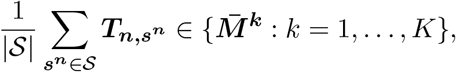

where 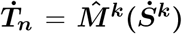 if *x*_*n,k*_ = 1, that is, antibiotic *k* is applied in step *n*. In order to test the success of our solution, we adopt an out-of-sample evaluation scheme. In particular, we generate a fresh sample of 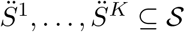 and evaluate the objective function as

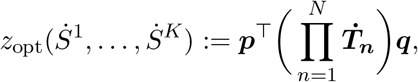

Where 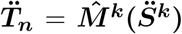. Our computational experiments in Section 3 reveal that choosing 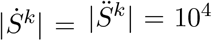 gives quite satisfactory results since the values *z*_opt_ and *z*_ev_ are typically very close to each other (recall that we have |*𝒮*| = *R*^*d*^ = 12^16^ in the data acquired from [21]).

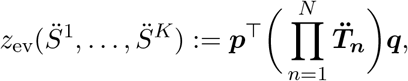

#### 2.2.3 Dynamic Optimization

In this section, we will consider the dynamic optimization of the Antibiotics Time Machine Problem in the sense that the treatment plan is decided throughout the planning horizon as the state transitions are observed.

The following proposition, whose proof is similar to that of Proposition 2, is a key result.

##### Proposition 3.

*Problem 4 is an instance of Problem 2*.

If the set of average transition probability matrices 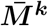 are available, then we can directly use the solution approaches from Section 2.1.4. However, this is typically not the case and due to the discussion from the previous section, we will resort to sampling. In particular, given samples 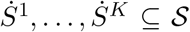, we first compute the set of estimator matrices 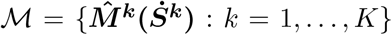 using equation (10). We then use the DP approach from Subsection 2.1.4 to obtain a policy, which we will denote by *y*_*n,i*_. Let us denote the optimal value with respect to the sample 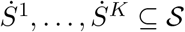 as *z*_opt_ again.

To estimate the objective function value of the policy using an out-of-sample analysis, we generate a fresh sample of 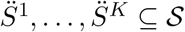 and compute the estimator matrices as 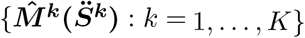. We use a forward DP approach given in Algorithm 2 to evaluate the objective function value of the policy with respect to the sample 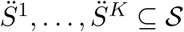, which we will denote as *z*_ev_ again.

##### Algorithm 2 Forward DP.

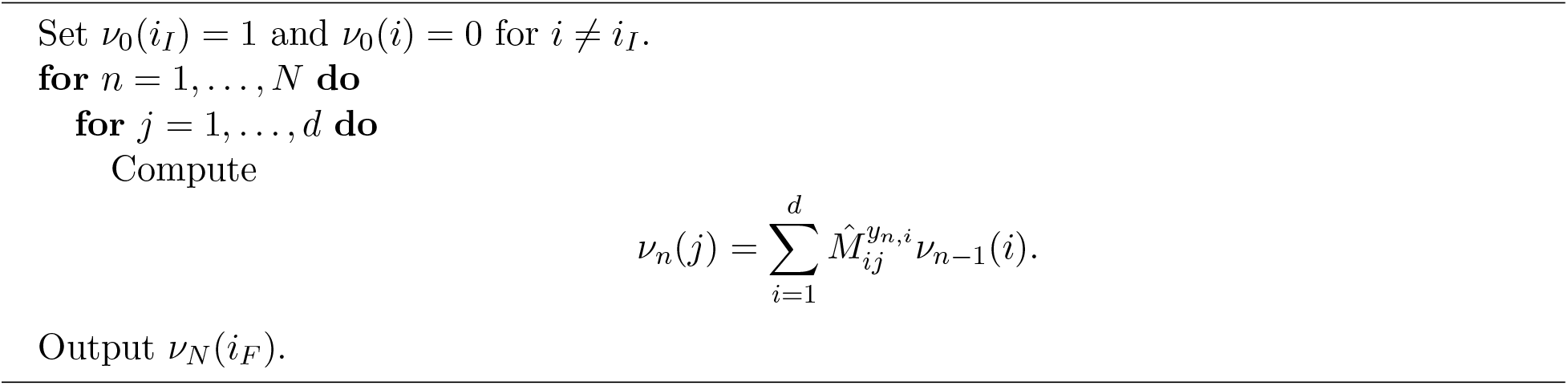

We note that the running time of Algorithm 2 is *O*(*d*^2^*N*).

## 3 Results and Discussion

### 3.1 Computational Setting

We use the experimental growth rate data from [21] to obtain the transition probability matrices. This dataset consists of the growth rates of *d* = 16 genotypes under *K* = 23 different antibiotics (or doses) measured for *R* = 12 times (see Table 1). We select 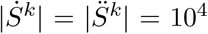 to estimate the average transition probability matrices for *k* = 1, …, *K*. We add the *d × d* identity matrix as the zeroth antibiotic matrix to model the “no intake” action.

**Table 1:** List of antibiotics and doses (in μg*/*mL) from [21].

| Code | Antibiotic | Doses | Code | Antibiotic | Doses |
| --- | --- | --- | --- | --- | --- |
| AM | Amoxicillin | 256, 512 | CRO | Ceftriaxone | 0.025, 0.5, 1 |
| AMC | AM/Clavulanate | 8/8 | CXM | Cefuroxime | 2.25, 3, 4 |
| CAZ | Ceftazidime | 0.125, 0.25, 0.5 | FEP | Cefepime | 0.0312, 0.0625, 0.125 |
| CEC | Cefaclor | 4 | TZP | Piperacillin/Tazobactam | 8/32, 8/64, 8/128 |
| CPR | Cefprozil | 8, 12, 16 | ZOX | Ceftizoxime | 0.0156 |

We conduct our computational experiments on a 64-bit workstation with two Intel(R) Xeon(R) Gold 6248R CPU (3.00GHz) processors (256 GB RAM) using the Python programming language. We utilize Gurobi 9.5 to solve the MILPs with the default settings (except for the absolute optimality gap parameter, which is set to 0.001, and the time limit, which is set to one hour).

To reproduce and verify computational results, we have uploaded the source code to GitHub repository [27]. Detailed instructions can be found under README.md of this repository.

We also provide an online companion entitled Detailed Results.xlsx, which includes the results of all experiments up to *N* = 16 in terms of the objective function values and computational times in seconds.

### 3.2 Results of the Static Stochastic Version

In this section, we present the results of our computational experiments for the Static Stochastic Version of the Antibiotics Time Machine Problem, that is, Problem 3. In Table 2 (resp. Table 3), we report the SAA results under CPM (resp. EPM) for different *N* values. In particular, for each initial genotype *i*_*I*_, we tabulate the objective function value of reaching the wild type in at most *N* steps obtained from the MILP 5 as *z*_opt_, and the evaluated objective value from out-of-sample as *z*_ev_. In addition, we give the average CPU time (in seconds) of the MILP (5) model as “AT(s)” in the last row.

**Table 2:** SAA Results of Static Stochastic Version under CPM.

| | $N = 1$ | | $N = 2$ | | $N = 3$ | | $N = 4$ | | $N = 5$ | | $N = 6$ | | $N = 9$ | | $N = 12$ | | $N = 16$ | |
| --- | --- | --- | --- | --- | --- | --- | --- | --- | --- | --- | --- | --- | --- | --- | --- | --- | --- | --- |
| $i_I$ | $z_{\text{opt}}$ | $z_{\text{ev}}$ | $z_{\text{opt}}$ | $z_{\text{ev}}$ | $z_{\text{opt}}$ | $z_{\text{ev}}$ | $z_{\text{opt}}$ | $z_{\text{ev}}$ | $z_{\text{opt}}$ | $z_{\text{ev}}$ | $z_{\text{opt}}$ | $z_{\text{ev}}$ | $z_{\text{opt}}$ | $z_{\text{ev}}$ | $z_{\text{opt}}$ | $z_{\text{ev}}$ | $z_{\text{opt}}$ | $z_{\text{ev}}$ |
| 0001 | 0.46 | 0.46 | 0.46 | 0.46 | 0.49 | 0.49 | 0.49 | 0.49 | 0.54 | 0.54 | 0.58 | 0.58 | 0.69 | 0.69 | 0.71 | 0.71 | 0.73 | 0.73 |
| 0010 | 0.65 | 0.65 | 0.65 | 0.65 | 0.77 | 0.77 | 0.77 | 0.77 | 0.77 | 0.77 | 0.77 | 0.77 | 0.77 | 0.77 | 0.77 | 0.77 | 0.77 | 0.77 |
| 0011 | 0.00 | 0.00 | 0.58 | 0.58 | 0.58 | 0.58 | 0.72 | 0.72 | 0.72 | 0.72 | 0.74 | 0.74 | 0.74 | 0.74 | 0.75 | 0.75 | 0.75 | 0.75 |
| 0100 | 0.94 | 0.94 | 0.94 | 0.94 | 0.94 | 0.94 | 0.94 | 0.94 | 0.94 | 0.94 | 0.94 | 0.94 | 0.94 | 0.94 | 0.94 | 0.94 | 0.94 | 0.94 |
| 0101 | 0.00 | 0.00 | 0.45 | 0.46 | 0.45 | 0.46 | 0.54 | 0.54 | 0.56 | 0.56 | 0.66 | 0.67 | 0.70 | 0.70 | 0.72 | 0.72 | 0.73 | 0.73 |
| 0110 | 0.00 | 0.00 | 0.77 | 0.77 | 0.77 | 0.77 | 0.77 | 0.77 | 0.77 | 0.77 | 0.77 | 0.77 | 0.77 | 0.77 | 0.77 | 0.77 | 0.77 | 0.77 |
| 0111 | 0.00 | 0.00 | 0.00 | 0.00 | 0.59 | 0.59 | 0.61 | 0.61 | 0.71 | 0.71 | 0.71 | 0.71 | 0.73 | 0.73 | 0.74 | 0.74 | 0.74 | 0.75 |
| 1000 | 0.62 | 0.62 | 0.63 | 0.63 | 0.63 | 0.63 | 0.63 | 0.63 | 0.63 | 0.63 | 0.63 | 0.63 | 0.63 | 0.63 | 0.67 | 0.67 | 0.70 | 0.71 |
| 1001 | 0.00 | 0.00 | 0.52 | 0.52 | 0.57 | 0.57 | 0.57 | 0.57 | 0.57 | 0.57 | 0.57 | 0.57 | 0.59 | 0.59 | 0.65 | 0.66 | 0.70 | 0.70 |
| 1010 | 0.00 | 0.00 | 0.60 | 0.60 | 0.60 | 0.60 | 0.70 | 0.70 | 0.70 | 0.70 | 0.70 | 0.70 | 0.71 | 0.71 | 0.72 | 0.72 | 0.73 | 0.73 |
| 1011 | 0.00 | 0.00 | 0.00 | 0.00 | 0.44 | 0.44 | 0.48 | 0.48 | 0.54 | 0.54 | 0.62 | 0.62 | 0.67 | 0.67 | 0.70 | 0.70 | 0.72 | 0.73 |
| 1100 | 0.00 | 0.00 | 0.59 | 0.59 | 0.60 | 0.60 | 0.60 | 0.60 | 0.60 | 0.60 | 0.60 | 0.60 | 0.60 | 0.60 | 0.66 | 0.66 | 0.70 | 0.71 |
| 1101 | 0.00 | 0.00 | 0.00 | 0.00 | 0.52 | 0.52 | 0.57 | 0.57 | 0.57 | 0.57 | 0.57 | 0.57 | 0.59 | 0.59 | 0.66 | 0.66 | 0.70 | 0.70 |
| 1110 | 0.00 | 0.00 | 0.00 | 0.00 | 0.64 | 0.64 | 0.64 | 0.64 | 0.65 | 0.65 | 0.65 | 0.65 | 0.69 | 0.69 | 0.71 | 0.71 | 0.73 | 0.73 |
| 1111 | 0.00 | 0.00 | 0.00 | 0.00 | 0.00 | 0.00 | 0.52 | 0.52 | 0.57 | 0.56 | 0.57 | 0.56 | 0.62 | 0.62 | 0.66 | 0.67 | 0.70 | 0.70 |
| AT(s) | 0.003 |  | 0.018 |  | 0.088 |  | 0.296 |  | 1.214 |  | 2.443 |  | 16.880 |  | 168.888 |  | 2655.636 |  |

**Table 3:** SAA Results of Static Stochastic Version under EPM.

| | $N = 1$ | | $N = 2$ | | $N = 3$ | | $N = 4$ | | $N = 5$ | | $N = 6$ | | $N = 9$ | | $N = 12$ | | $N = 16$ | |
| --- | --- | --- | --- | --- | --- | --- | --- | --- | --- | --- | --- | --- | --- | --- | --- | --- | --- | --- |
| $i_I$ | $z_{\text{opt}}$ | $z_{\text{ev}}$ | $z_{\text{opt}}$ | $z_{\text{ev}}$ | $z_{\text{opt}}$ | $z_{\text{ev}}$ | $z_{\text{opt}}$ | $z_{\text{ev}}$ | $z_{\text{opt}}$ | $z_{\text{ev}}$ | $z_{\text{opt}}$ | $z_{\text{ev}}$ | $z_{\text{opt}}$ | $z_{\text{ev}}$ | $z_{\text{opt}}$ | $z_{\text{ev}}$ | $z_{\text{opt}}$ | $z_{\text{ev}}$ |
| 0001 | 0.45 | 0.45 | 0.45 | 0.45 | 0.45 | 0.45 | 0.46 | 0.46 | 0.47 | 0.47 | 0.48 | 0.48 | 0.49 | 0.49 | 0.49 | 0.49 | 0.50 | 0.50 |
| 0010 | 0.57 | 0.57 | 0.57 | 0.57 | 0.57 | 0.57 | 0.57 | 0.57 | 0.57 | 0.57 | 0.57 | 0.57 | 0.57 | 0.57 | 0.57 | 0.57 | 0.57 | 0.57 |
| 0011 | 0.00 | 0.00 | 0.46 | 0.46 | 0.46 | 0.46 | 0.47 | 0.47 | 0.47 | 0.47 | 0.47 | 0.48 | 0.49 | 0.48 | 0.49 | 0.49 | 0.50 | 0.50 |
| 0100 | 0.51 | 0.50 | 0.51 | 0.50 | 0.51 | 0.50 | 0.51 | 0.50 | 0.51 | 0.50 | 0.51 | 0.50 | 0.51 | 0.50 | 0.51 | 0.50 | 0.51 | 0.50 |
| 0101 | 0.00 | 0.00 | 0.39 | 0.40 | 0.39 | 0.40 | 0.42 | 0.42 | 0.46 | 0.46 | 0.48 | 0.48 | 0.50 | 0.50 | 0.50 | 0.50 | 0.51 | 0.50 |
| 0110 | 0.00 | 0.00 | 0.52 | 0.52 | 0.52 | 0.52 | 0.53 | 0.53 | 0.53 | 0.53 | 0.53 | 0.53 | 0.53 | 0.53 | 0.53 | 0.53 | 0.53 | 0.53 |
| 0111 | 0.00 | 0.00 | 0.00 | 0.00 | 0.39 | 0.39 | 0.41 | 0.40 | 0.42 | 0.42 | 0.45 | 0.44 | 0.49 | 0.49 | 0.50 | 0.49 | 0.50 | 0.50 |
| 1000 | 0.60 | 0.60 | 0.62 | 0.62 | 0.62 | 0.62 | 0.62 | 0.62 | 0.62 | 0.62 | 0.62 | 0.62 | 0.62 | 0.62 | 0.62 | 0.62 | 0.62 | 0.62 |
| 1001 | 0.00 | 0.00 | 0.50 | 0.50 | 0.54 | 0.54 | 0.54 | 0.54 | 0.54 | 0.54 | 0.54 | 0.54 | 0.54 | 0.54 | 0.54 | 0.54 | 0.54 | 0.54 |
| 1010 | 0.00 | 0.00 | 0.47 | 0.48 | 0.47 | 0.48 | 0.47 | 0.48 | 0.47 | 0.48 | 0.48 | 0.47 | 0.49 | 0.49 | 0.49 | 0.49 | 0.50 | 0.50 |
| 1011 | 0.00 | 0.00 | 0.00 | 0.00 | 0.37 | 0.37 | 0.39 | 0.39 | 0.43 | 0.42 | 0.47 | 0.47 | 0.49 | 0.49 | 0.50 | 0.49 | 0.50 | 0.50 |
| 1100 | 0.00 | 0.00 | 0.36 | 0.36 | 0.36 | 0.36 | 0.49 | 0.49 | 0.53 | 0.53 | 0.53 | 0.53 | 0.53 | 0.53 | 0.53 | 0.53 | 0.53 | 0.53 |
| 1101 | 0.00 | 0.00 | 0.00 | 0.00 | 0.50 | 0.50 | 0.54 | 0.54 | 0.54 | 0.54 | 0.54 | 0.54 | 0.54 | 0.54 | 0.54 | 0.54 | 0.54 | 0.54 |
| 1110 | 0.00 | 0.00 | 0.00 | 0.00 | 0.30 | 0.30 | 0.31 | 0.31 | 0.46 | 0.46 | 0.49 | 0.49 | 0.50 | 0.50 | 0.50 | 0.50 | 0.51 | 0.50 |
| 1111 | 0.00 | 0.00 | 0.00 | 0.00 | 0.00 | 0.00 | 0.44 | 0.44 | 0.47 | 0.47 | 0.50 | 0.49 | 0.51 | 0.51 | 0.52 | 0.51 | 0.52 | 0.51 |
| AT(s) | 0.010 |  | 0.018 |  | 0.082 |  | 0.320 |  | 1.207 |  | 2.136 |  | 10.465 |  | 90.005 |  | 1803.956 |  |

We have several observations derived from these experiments: Firstly, the values *z*_opt_ and *z*_ev_ are very close to each other, which justifies the accuracy of our SAA-based approach. In fact, we have |*z*_opt_ *− z*_ev_| *≤* 0.01 in all cases. Secondly, the objective values increase as *N* gets larger, in general. Under CPM (resp. EPM), the probability of reaching the wild type exceeds 0.695 (resp. 0.495) for all initial genotypes for *N* = 16. Lastly, the average CPU time increases sharply as *N* becomes larger. We also observe that the CPU time under CPM is typically longer than the CPU time under EPM, and the solver reaches the one-hour time limit frequently for *N ≥* 14 under CPM.

### 3.3 Results of the Dynamic Stochastic Version

In this section, we present the results of our computational experiments for the Dynamic Stochastic Version of the Antibiotics Time Machine Problem, that is, Problem 4. In Table 4 (resp. Table 5), we report the SAA results under CPM (resp. EPM) for different *N* values. In particular, for each initial genotype *i*_*I*_, we tabulate the objective function value of reaching the wild type in at most *N* steps obtained from the backward DP Algorithm 1 as *z*_opt_, and the evaluated objective value from out-of-sample obtained from the forward DP Algorithm 2 as *z*_ev_. Since the DP algorithms are quite fast, we do not report their CPU times, which are less than a second.

**Table 4:** SAA Results of Dynamic Stochastic Version under CPM.

| | $N = 1$ | | $N = 2$ | | $N = 3$ | | $N = 4$ | | $N = 5$ | | $N = 6$ | | $N = 9$ | | $N = 11$ | |
| --- | --- | --- | --- | --- | --- | --- | --- | --- | --- | --- | --- | --- | --- | --- | --- | --- |
| $i_I$ | $z_{\text{opt}}$ | $z_{\text{ev}}$ | $z_{\text{opt}}$ | $z_{\text{ev}}$ | $z_{\text{opt}}$ | $z_{\text{ev}}$ | $z_{\text{opt}}$ | $z_{\text{ev}}$ | $z_{\text{opt}}$ | $z_{\text{ev}}$ | $z_{\text{opt}}$ | $z_{\text{ev}}$ | $z_{\text{opt}}$ | $z_{\text{ev}}$ | $z_{\text{opt}}$ | $z_{\text{ev}}$ |
| 0001 | 0.46 | 0.46 | 0.57 | 0.57 | 0.77 | 0.77 | 0.86 | 0.86 | 0.93 | 0.94 | 0.96 | 0.96 | 1.00 | 1.00 | 1.00 | 1.00 |
| 0010 | 0.65 | 0.65 | 0.80 | 0.81 | 0.93 | 0.93 | 0.95 | 0.95 | 0.98 | 0.98 | 0.99 | 0.99 | 1.00 | 1.00 | 1.00 | 1.00 |
| 0011 | 0.00 | 0.00 | 0.58 | 0.58 | 0.71 | 0.72 | 0.90 | 0.90 | 0.93 | 0.93 | 0.97 | 0.97 | 1.00 | 1.00 | 1.00 | 1.00 |
| 0100 | 0.94 | 0.94 | 0.94 | 0.94 | 0.97 | 0.97 | 0.98 | 0.98 | 0.99 | 0.99 | 0.99 | 0.99 | 1.00 | 1.00 | 1.00 | 1.00 |
| 0101 | 0.00 | 0.00 | 0.45 | 0.46 | 0.57 | 0.57 | 0.77 | 0.77 | 0.86 | 0.86 | 0.93 | 0.93 | 0.99 | 0.99 | 1.00 | 1.00 |
| 0110 | 0.00 | 0.00 | 0.80 | 0.80 | 0.84 | 0.85 | 0.94 | 0.94 | 0.96 | 0.96 | 0.98 | 0.98 | 1.00 | 1.00 | 1.00 | 1.00 |
| 0111 | 0.00 | 0.00 | 0.00 | 0.00 | 0.70 | 0.70 | 0.77 | 0.78 | 0.91 | 0.91 | 0.94 | 0.94 | 0.99 | 0.99 | 1.00 | 1.00 |
| 1000 | 0.62 | 0.62 | 0.80 | 0.80 | 0.90 | 0.90 | 0.95 | 0.95 | 0.97 | 0.97 | 0.99 | 0.99 | 1.00 | 1.00 | 1.00 | 1.00 |
| 1001 | 0.00 | 0.00 | 0.53 | 0.53 | 0.73 | 0.73 | 0.86 | 0.86 | 0.92 | 0.92 | 0.96 | 0.96 | 0.99 | 0.99 | 1.00 | 1.00 |
| 1010 | 0.00 | 0.00 | 0.63 | 0.63 | 0.79 | 0.80 | 0.91 | 0.91 | 0.95 | 0.95 | 0.98 | 0.97 | 1.00 | 1.00 | 1.00 | 1.00 |
| 1011 | 0.00 | 0.00 | 0.00 | 0.00 | 0.55 | 0.55 | 0.72 | 0.72 | 0.88 | 0.88 | 0.93 | 0.93 | 0.99 | 0.99 | 1.00 | 1.00 |
| 1100 | 0.00 | 0.00 | 0.59 | 0.59 | 0.76 | 0.76 | 0.89 | 0.89 | 0.94 | 0.94 | 0.97 | 0.97 | 1.00 | 1.00 | 1.00 | 1.00 |
| 1101 | 0.00 | 0.00 | 0.00 | 0.00 | 0.53 | 0.53 | 0.73 | 0.73 | 0.86 | 0.86 | 0.92 | 0.92 | 0.99 | 0.99 | 1.00 | 1.00 |
| 1110 | 0.00 | 0.00 | 0.00 | 0.00 | 0.70 | 0.70 | 0.78 | 0.78 | 0.91 | 0.91 | 0.94 | 0.94 | 0.99 | 0.99 | 1.00 | 1.00 |
| 1111 | 0.00 | 0.00 | 0.00 | 0.00 | 0.00 | 0.00 | 0.62 | 0.62 | 0.75 | 0.75 | 0.89 | 0.89 | 0.98 | 0.98 | 1.00 | 1.00 |

**Table 5:** SAA Results of Dynamic Stochastic Version under EPM.

| | $N = 1$ | | $N = 2$ | | $N = 3$ | | $N = 4$ | | $N = 5$ | | $N = 6$ | | $N = 9$ | | $N = 12$ | | $N = 14$ | |
| --- | --- | --- | --- | --- | --- | --- | --- | --- | --- | --- | --- | --- | --- | --- | --- | --- | --- | --- |
| $i_I$ | $z_{\text{opt}}$ | $z_{\text{ev}}$ | $z_{\text{opt}}$ | $z_{\text{ev}}$ | $z_{\text{opt}}$ | $z_{\text{ev}}$ | $z_{\text{opt}}$ | $z_{\text{ev}}$ | $z_{\text{opt}}$ | $z_{\text{ev}}$ | $z_{\text{opt}}$ | $z_{\text{ev}}$ | $z_{\text{opt}}$ | $z_{\text{ev}}$ | $z_{\text{opt}}$ | $z_{\text{ev}}$ | $z_{\text{opt}}$ | $z_{\text{ev}}$ |
| 0001 | 0.45 | 0.45 | 0.55 | 0.55 | 0.72 | 0.72 | 0.82 | 0.82 | 0.89 | 0.89 | 0.93 | 0.93 | 0.98 | 0.98 | 1.00 | 1.00 | 1.00 | 1.00 |
| 0010 | 0.57 | 0.57 | 0.74 | 0.75 | 0.85 | 0.86 | 0.91 | 0.91 | 0.94 | 0.95 | 0.96 | 0.97 | 0.99 | 0.99 | 1.00 | 1.00 | 1.00 | 1.00 |
| 0011 | 0.00 | 0.00 | 0.46 | 0.46 | 0.62 | 0.63 | 0.77 | 0.77 | 0.85 | 0.86 | 0.91 | 0.91 | 0.98 | 0.98 | 0.99 | 1.00 | 1.00 | 1.00 |
| 0100 | 0.51 | 0.50 | 0.61 | 0.60 | 0.71 | 0.71 | 0.78 | 0.78 | 0.87 | 0.87 | 0.91 | 0.91 | 0.98 | 0.98 | 1.00 | 1.00 | 1.00 | 1.00 |
| 0101 | 0.00 | 0.00 | 0.40 | 0.40 | 0.49 | 0.49 | 0.69 | 0.69 | 0.80 | 0.80 | 0.88 | 0.88 | 0.97 | 0.97 | 0.99 | 0.99 | 1.00 | 1.00 |
| 0110 | 0.00 | 0.00 | 0.53 | 0.53 | 0.71 | 0.72 | 0.83 | 0.84 | 0.89 | 0.90 | 0.94 | 0.94 | 0.98 | 0.99 | 1.00 | 1.00 | 1.00 | 1.00 |
| 0111 | 0.00 | 0.00 | 0.00 | 0.00 | 0.45 | 0.45 | 0.61 | 0.61 | 0.77 | 0.77 | 0.85 | 0.86 | 0.97 | 0.97 | 0.99 | 0.99 | 1.00 | 1.00 |
| 1000 | 0.60 | 0.60 | 0.80 | 0.79 | 0.90 | 0.89 | 0.94 | 0.94 | 0.97 | 0.97 | 0.98 | 0.98 | 1.00 | 1.00 | 1.00 | 1.00 | 1.00 | 1.00 |
| 1001 | 0.00 | 0.00 | 0.50 | 0.51 | 0.70 | 0.70 | 0.83 | 0.83 | 0.90 | 0.90 | 0.94 | 0.94 | 0.99 | 0.99 | 1.00 | 1.00 | 1.00 | 1.00 |
| 1010 | 0.00 | 0.00 | 0.50 | 0.49 | 0.65 | 0.65 | 0.80 | 0.81 | 0.87 | 0.87 | 0.93 | 0.93 | 0.98 | 0.98 | 1.00 | 1.00 | 1.00 | 1.00 |
| 1011 | 0.00 | 0.00 | 0.00 | 0.00 | 0.43 | 0.43 | 0.59 | 0.59 | 0.77 | 0.77 | 0.85 | 0.85 | 0.97 | 0.97 | 0.99 | 0.99 | 1.00 | 1.00 |
| 1100 | 0.00 | 0.00 | 0.37 | 0.37 | 0.47 | 0.46 | 0.71 | 0.70 | 0.81 | 0.81 | 0.89 | 0.89 | 0.98 | 0.98 | 0.99 | 0.99 | 1.00 | 1.00 |
| 1101 | 0.00 | 0.00 | 0.00 | 0.00 | 0.50 | 0.50 | 0.70 | 0.70 | 0.83 | 0.83 | 0.90 | 0.90 | 0.98 | 0.98 | 1.00 | 1.00 | 1.00 | 1.00 |
| 1110 | 0.00 | 0.00 | 0.00 | 0.00 | 0.39 | 0.39 | 0.52 | 0.52 | 0.74 | 0.74 | 0.83 | 0.83 | 0.96 | 0.96 | 0.99 | 0.99 | 1.00 | 1.00 |
| 1111 | 0.00 | 0.00 | 0.00 | 0.00 | 0.00 | 0.00 | 0.48 | 0.48 | 0.67 | 0.67 | 0.81 | 0.81 | 0.96 | 0.96 | 0.99 | 0.99 | 1.00 | 1.00 |

We now discuss our observations derived from these experiments: Firstly, the values *z*_opt_ and *z*_ev_ are again very close to each other. Similar to the experiments in the previous subsection, we have |*z*_opt_ *− z*_ev_| *≤* 0.01 in all cases considered. Secondly, the expected probabilities of reaching the wild type with the Dynamic Version are strictly better than that with the Static Version, which justifies the added flexibility. In fact, the objective values exceed 0.995 for CPM (resp. EPM) when *N* = 11 (resp. *N* = 14), which are not attainable with the static versions.

### 3.4 Visualization of Optimal Solutions for Two Specific Cases

In this section, we present the optimal solutions of the Static and Dynamic Stochastic Versions of the Antibiotics Time Machine Problem for two specific cases to facilitate the discussion about their differences. We set the number of steps as *N* = 3 in an attempt to have simple enough illustrations with meaningful and insightful observations. The optimal solutions reported below use doses of 512, 0.1, 12 and 8/32 for AM, CRO, CPR and TZP, respectively (the other antibiotics involved have a single dose as reported in Table 1).

For the first illustration, we choose the initial genotype *i*_*I*_ = 1010 with the number of steps *N* = 3 under CPM, and visualize the optimal solutions in Figure 5. Here, we have *N* + 1 = 4 layers of nodes where circular-shaped nodes indicate antibiotic decisions taken in that genotype. The optimal sequence for the Static Version is CRO-AM-I (where I represents the no-intake action) while the optimal policy for the Dynamic Version is a function of the genotype and the step. For instance, the optimal policy states that we should apply CPR in Step 3 if the genotype is 1000. Diamond-shaped nodes give the probability of reaching that terminal genotype. For example, the probability of reaching the wild type is 0.597 and 0.792 for the Static and Dynamic Versions, respectively. Transition probabilities above 0.01 are shown as the corresponding arc labels between a pair of nodes while arcs with probabilities below this threshold are omitted in an effort not to overcomplicate the figures.

**Figure 5:**
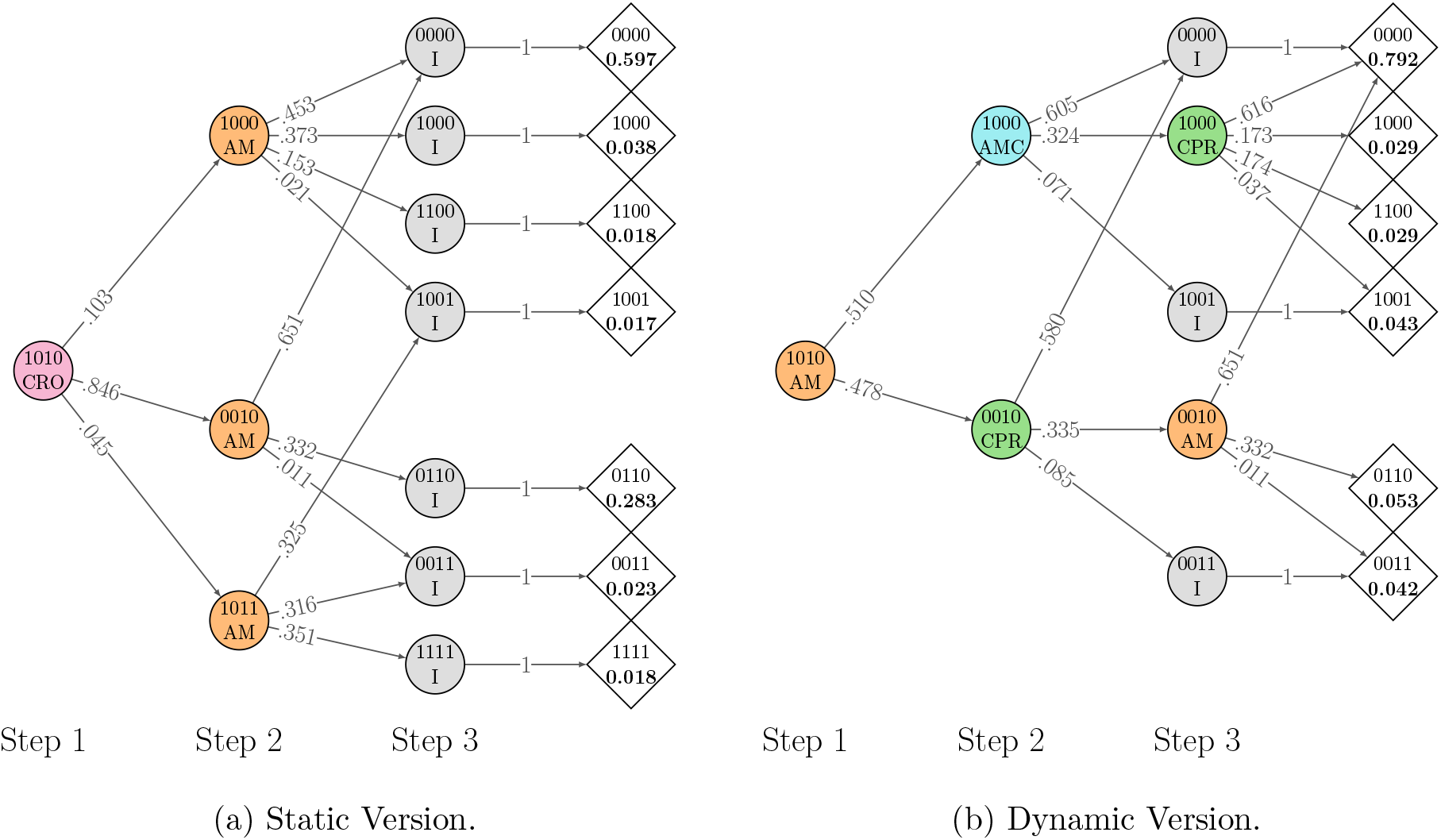
Visualization of optimal solutions for *i*_*I*_ = 1010 with *N* = 3 under CPM.

We have several interesting observations related to Figure 5. Firstly, the probability of reaching the wild type in the Dynamic Version is 0.792 (rounded as 0.79 in row 1010 and column *z*_opt_ of Table 4) whereas this probability is 0.597 for the Static Version (rounded as 0.60 in row 1010 and column *z*_opt_ of Table 2), which is significantly smaller. Recall from Table 2 that the probability goes up to only 0.73 for this initial genotype with the Static Version even when *N* = 16 while the Dynamic Version quickly reaches 0.98 when *N* = 6 according to Table 4. Secondly, the Static Version takes a no-intake action in Step 3, which means that this step is essentially “wasted”. This partly explains the first observation why the Static Version is much worse than the Dynamic Version. Thirdly, we observe that multiple paths might lead to the wild type. For example, there are respectively two and four such paths in the Static and Dynamic Versions, which again partly explains the first observation. The fourth and last observation is more subtle: The dynamic optimization is not “greedy”, meaning that it does not necessarily choose the antibiotic which has the largest probability of reaching the wild type in every step. For example, let us focus on genotype 1000. In Step 2, the optimization chooses AMC with the probability of reaching the wild type as 0.605 whereas in Step 3, the optimal decision is to apply CPR with the probability of 0.616. The reason is that if CPR were applied in Step 2, then with probability 0.211, the next genotype would have two mutations, which is impossible to reverse in one step. This probability is much smaller for AMC, making it a less riskier choice in Step 2. On the other hand, it is optimal to apply CPR in Step 3 as this antibiotic has the largest probability of reaching the wild type in one step for genotype 1000. A similar situation is observed for the genotype 0010 as well.

For the second illustration, we choose the initial genotype *i*_*I*_ = 1011 with the number of steps *N* = 3 under CPM, and visualize the optimal solutions in Figure 6 with the same notation as in the previous illustration. Again, we have some interesting observations. Firstly, the probability of reaching the wild type in the Dynamic Version is 0.552 (rounded as 0.55 in row 1011 and column *z*_opt_ of Table 4) whereas this probability is 0.437 for the Static Version (rounded as 0.44 in row 1011 and column *z*_opt_ of Table 2). Secondly, unlike the previous illustration, no action is wasted in the static version, which is an expected behavior since it is impossible to reach the wild type otherwise. This also partly explains the relatively small difference in terms of the optimal values of static and dynamic versions. Thirdly, compared to Figure 5 related to the first illustration, we observe denser networks in Figure 6. This is due to the fact that the initial genotype has more mutations, which diversifies the possible pathways.

**Figure 6:**
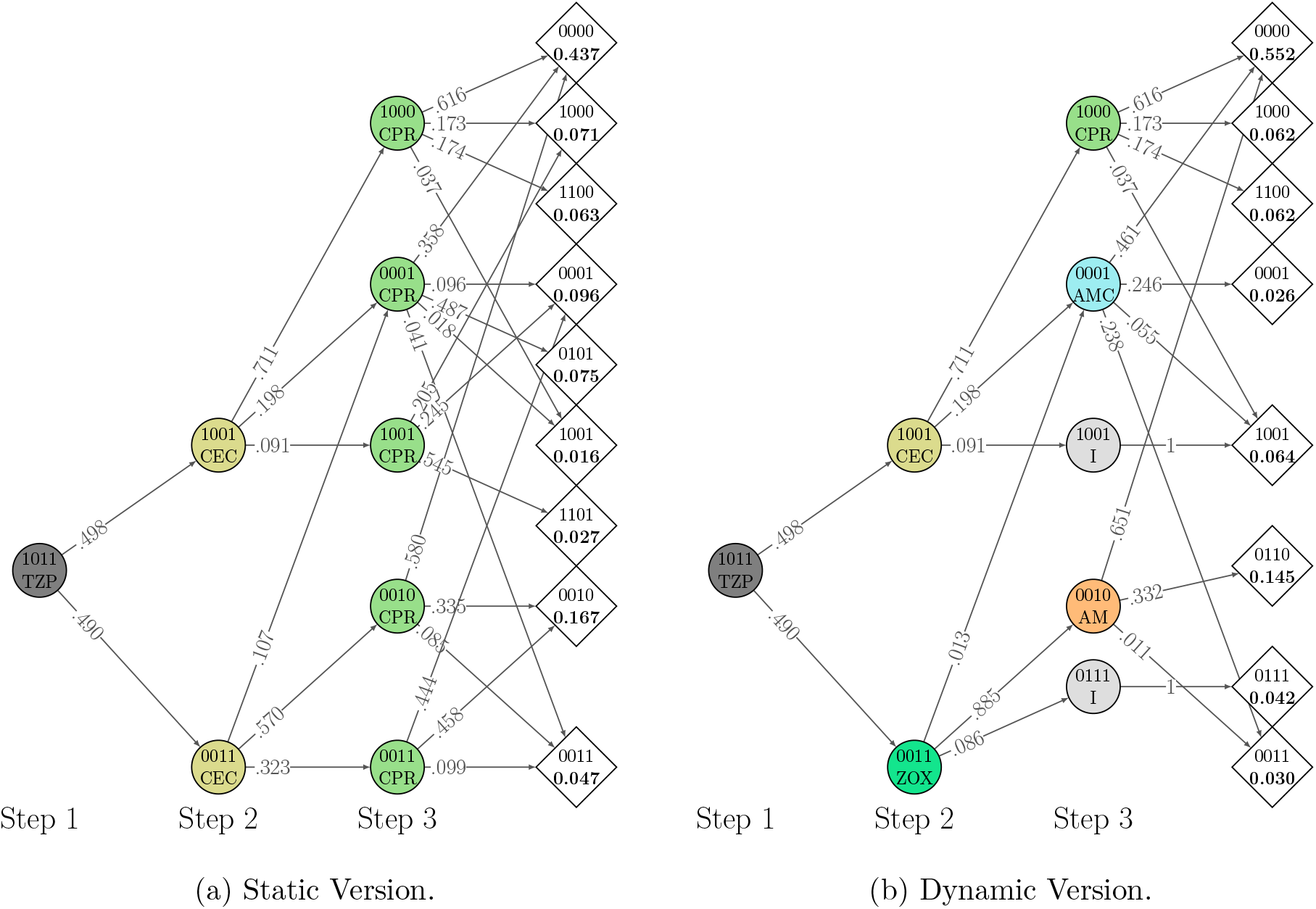
Visualization of optimal solutions for *i*_*I*_ = 1011 with *N* = 3 under CPM.

Note that the distinguishing factor of the above illustrations is the initial genotypes, which are respectively chosen as 1010 and 1011. We have seen that the number of mutations in the initial genotypes lead to different observations, especially when the number of steps is chosen as small. Finally, a property common in both illustrations is worth mentioning: The static versions lead to denser networks compared to the dynamic versions since the latter is more informed due to the intermediate observations and make more targeted decisions towards reaching the wild type in general.

## 4 Conclusions

We have developed stochastic dynamic programming approach to recommend an adaptive drug delivery policy. We have provided toy examples to make a distinction between preset (static) and adaptive (dynamic) as well as between deterministic (single measurement for each growth rate) and stochastic (multiple measurements) approaches. We conduct extensive computational experiments on the four-allele landscape of the *E. coli β*-lactamase gene for static-stochastic and dynamic-stochastic cases. This is done for 23 different drugs/doses and the results are compared. The success of the adaptive methodology for multiple measurements has shown to be remarkable. We encourage laboratory and if possible clinic studies to test the efficacy of our suggested policies.

## Supporting information

Detailed Results

## Footnotes

1 With some abuse of notation, we will interchangeably use the terms genotype and its ID for convenience.

## References

[1] Murray CJ, Ikuta KS, Sharara F, Swetschinski L, Aguilar GR, Gray A, et al. Global burden of bacterial antimicrobial resistance in 2019: a systematic analysis. The Lancet. 2022;399(10325):629–655.

[2] May M. How to fight antibiotic resistance. Nature Medicine. 2023;29(7):1583–1586.

[3] Liu G, Catacutan DB, Rathod K, Swanson K, Jin W, Mohammed JC, et al. Deep learning-guided discovery of an antibiotic targeting Acinetobacter baumannii. Nature Chemical Biology. 2023:1–9.

[4] Wong F, de la Fuente-Nunez C, Collins JJ. Leveraging artificial intelligence in the fight against infectious diseases. Science. 2023;381(6654):164–170.

[5] Brown D. Antibiotic resistance breakers: can repurposed drugs fill the antibiotic discovery void? Nature reviews Drug discovery. 2015;14(12):821–832.

[6] Tyers M, Wright GD. Drug combinations: a strategy to extend the life of antibiotics in the 21st century. Nature Reviews Microbiology. 2019;17(3):141–155.

[7] Mira PM, Crona K, Greene D, Meza JC, Sturmfels B, Barlow M. Rational design of antibiotic treatment plans: a treatment strategy for managing evolution and reversing resistance. PloS one. 2015;10(5):e0122283.

[8] Nichol D, Jeavons P, Fletcher AG, Bonomo RA, Maini PK, Paul JL, et al. Steering evolution with sequential therapy to prevent the emergence of bacterial antibiotic resistance. PLoS computational biology. 2015;11(9):e1004493.

[9] Bergstrom CT, Lo M, Lipsitch M. Ecological theory suggests that antimicrobial cycling will not reduce antimicrobial resistance in hospitals. Proceedings of the National Academy of Sciences. 2004;101(36):13285–13290.

[10] van Duijn PJ, Verbrugghe W, Jorens PG, Spöhr F, Schedler D, Deja M, et al. The effects of antibiotic cycling and mixing on antibiotic resistance in intensive care units: a cluster-randomised crossover trial. The Lancet Infectious Diseases. 2018;18(4):401–409.

[11] Nichol D, Bonomo RA, Scott JG. It’s too soon to pull the plug on antibiotic cycling. The Lancet Infectious Diseases. 2018;18(5):493.

[12] Nichol D, Rutter J, Bryant C, Hujer AM, Lek S, Adams MD, et al. Antibiotic collateral sensitivity is contingent on the repeatability of evolution. Nature communications. 2019;10(1):334.

[13] Maltas J, Wood KB. Pervasive and diverse collateral sensitivity profiles inform optimal strategies to limit antibiotic resistance. PLoS biology. 2019;17(10):e3000515.

[14] Maltas J, Huynh A, Wood KB. Dynamic collateral sensitivity profiles highlight challenges and opportunities for optimizing antibiotic sequences. bioRxiv. 2021:2021–12.

[15] Aulin LB, Liakopoulos A, van der Graaf PH, Rozen DE, van Hasselt JC. Design principles of collateral sensitivity-based dosing strategies. Nature communications. 2021;12(1):5691.

[16] Batra A, Roemhild R, Rousseau E, Franzenburg S, Niemann S, Schulenburg H. High potency of sequential therapy with only β-lactam antibiotics. Elife. 2021;10:e68876.

[17] Kocuk B. Optimization Problems Involving Matrix Multiplication with Applications in Material Science and Biology. Engineering Optimization. 2022;54(5):786–804.

[18] Weaver DT, Maltas J, Scott JG. Reinforcement Learning informs optimal treatment strategies to limit antibiotic resistance. bioRxiv. 2023:2023–01.

[19] Tran NM, Yang J. Antibiotics Time Machines Are Hard to Build. Notices of the AMS. 2017;64(10):1136–1140.

[20] Gluzman M, Scott JG, Vladimirsky A. Optimizing adaptive cancer therapy: dynamic programming and evolutionary game theory. Proceedings of the Royal Society B. 2020;287(1925):20192454.

[21] Mira PM, Barlow M, Meza JC, Hall BG. Statistical Package for Growth Rates Made Easy. Molecular biology and evolution. 2017;34(12):3303–3309.

[22] Gillespie JH. A simple stochastic gene substitution model. Theoretical population biology. 1983;23(2):202–215.

[23] Gillespie JH. Molecular evolution over the mutational landscape. Evolution. 1984;38(5):1116–1129.

[24] Kleywegt AJ, Shapiro A, Homem-de Mello T. The sample average approximation method for stochastic discrete optimization. SIAM Journal on Optimization. 2002;12(2):479–502.

[25] Birge JR, Louveaux F. Introduction to stochastic programming. Springer Science & Business Media; 2011.

[26] Shapiro A, Dentcheva D, Ruszczy ski A. Lectures on stochastic programming: modeling and theory. SIAM; 2014.

[27] Mesum O, Kocuk B. Stochastic Antibiotic Code; 2023. https://github.com/oguzmes/StochasticAntibiotic.

